# Multiscale Probabilistic Modeling: A Bayesian Approach to Augment Mechanistic Models of Cell Signaling with Machine-Learning Predictions of Binding Affinity

**DOI:** 10.1101/2025.05.23.655795

**Authors:** Holly A. Huber, Stacey D. Finley

**Author notes:** Corresponding Author: Stacey D. Finley.

## Abstract

Computational models in systems biology are often underdetermined—that is, there is little data relative to the complexity and size of the model. The lack of data is primarily due to limits in our ability to observe specific biological systems and restricts the utility of computational models. However, there are a growing number of experimental databases in biology. While these databases provide more observations, they often do not have observations that match the system of interest exactly. For example, database measurements might be collected at different experimental conditions or on a different scale compared to the system of interest. Here, we investigate what information can be gleaned from generalizing databases across these differences in the context of modeling a specific system – cell signaling. Ultimately, our goal is to better determine models of specific systems, thereby increasing their utility. To do this, we propose a novel, multiscale, probabilistic framework. We use this framework to integrate measurements of protein structure from the Protein Data Bank and measurements of amino acid sequence from the Universal Protein Resource into the parameter inference of cell signaling models. Then, we quantify exactly what information is gained from these measurements when modeling cell signaling. We choose to investigate the utility of these databases in the context of dynamic cell signaling models because experimental measurements of the variables of interest, protein dynamics, are still quite limited. We find that we can successfully integrate measurements from these databases to significantly improve parameter estimation of signaling models. The impact of sequence and structure measurements on model predictions depends on the sensitivity of the prediction to perturbations in the parameter values. Overall, this study demonstrates that measurements of protein structure and amino acid sequence can be leveraged to better inform parameters in models of cell signaling.

**Author Summary:** Computational models of cell signaling have provided mechanistic insights into complex biological systems, including in physiological and disease settings. Accurate and predictive modeling critically depends on the precise estimation of model parameters, which is often hindered by the limited availability of experimental data. In this study, we present a novel multiscale probabilistic inference framework that broadens the scope of data types that can be leveraged for parameter estimation for models of cell signaling. The framework integrates a machine learning pipeline with a generalizable parameter inference approach, enabling the use of experimental data across scales. Specifically, we demonstrate that incorporating protein amino acid sequence and 3D structural data enhances parameter estimation compared to traditional measurements such as protein concentrations over time. Improving parameter estimation increases the robustness and applicability of cell signaling models. Ultimately, our framework facilitates use of a broader range of data and supports the development of predictive computational models that increase our understanding of cell signaling.

## 1 Introduction

The application of computational modeling to biological systems has generated a substantial surge in discoveries and insights into these systems; however, especially as systems become more complex, accurately determining computational models remains challenging (1). For example, it can be challenging to parameterize systems biology models, as we often do not have direct measurements of model parameters. Instead, parameters must be inferred using noisy, partial observations of the system of interest. For instance, a system of ordinary differential equations (ODEs) may be used to model intracellular signaling (2). Such mechanistic models are useful, having been applied to optimize CAR-T cell therapies or elucidate control mechanisms of cellular responses (3,4). However, we rarely have observations of all the model parameters—such as binding affinities or protein shuttling rates. Instead, these parameters must be inferred using observations of the model variables: protein concentrations over time. Timeseries measurements of protein concentrations are often sparse, limited to relative concentrations, and noisy. When such measurements are used for parameter estimation, these limitations can result in significant uncertainty in the model predictions, which limits the applicability of the model (5).

To overcome this limitation, in this work we propose a novel, multiscale, probabilistic modeling framework that expands the experimental measurements that may be integrated into the parameter estimation of cell signaling models. Figure 1 illustrates this framework. We hypothesize that despite differences in scale, measurements of protein structure and amino acid sequence can still facilitate the parameter inference of ODE models of cell signaling. To overcome this difference in scale, our framework includes a user-friendly machine learning (ML) pipeline that predicts binding affinity parameters of cell signaling models using sequence or structure. To integrate these affinity parameter predictions into the estimation of signaling models, our framework also includes a generalizable Bayesian inference approach. This probabilistic approach to parameter estimation allows us to formally quantify the information gained from augmenting traditional experimental measurements for parameter estimation of cell signaling modes, such as protein concentration dynamics, with measurements of sequence and structure.

**Figure 1.**
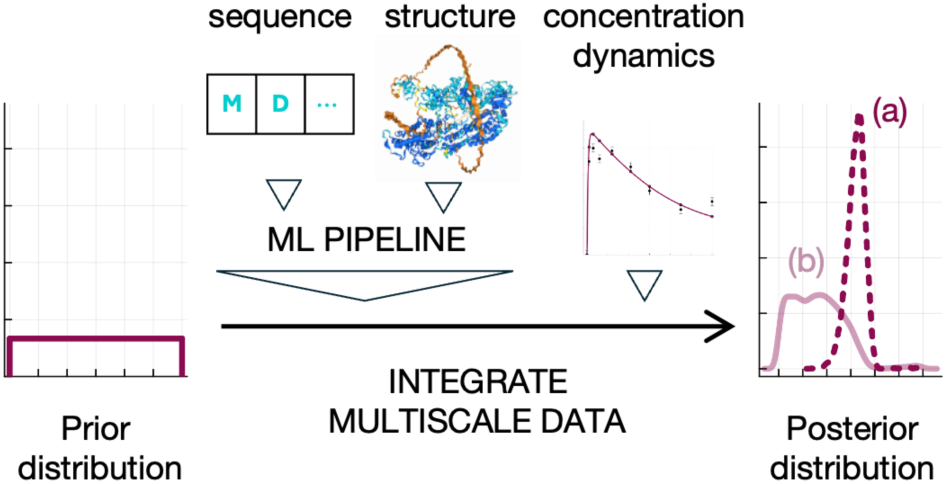
Multiscale Probabilistic Modeling Framework. Starting with the same range of possible parameter values—prior distribution—for the ODE model of cell signaling, we integrate data from different scales into our Bayesian parameter inference using a ML pipeline. We perform inference using two data sets. One dataset comprises only concentration dynamics. The other comprises concentration dynamics as well as amino acid sequence and protein structure. We may then compare the difference between the plausible parameter values—posterior distribution—with, (a), and without, (b), measurements of sequence and structure.

We use measurements of protein structure and amino acid sequence to augment our cell signaling model parameter inference. While experimental observations are still limited for the parameters and variables of signaling models, there have been significant breakthroughs in the ability to measure other biological variables. For example, researchers have been able to collect many more measurements of protein structure and amino acid sequence. This has resulted in rich databases such as the Protein Data Bank (PDB) and Universal Protein Resource (UniProt) (6,7). Thus, we propose augmenting timeseries measurements of protein concentrations with more plentiful data on sequence and structure.

Our proposed multiscale, probabilistic framework leverages a ML pipeline to integrate measurements of sequence and structure into the parameter inference of cell signaling models. Integrating such data into the parameter estimation is non-trivial because the data are not observations of variables or parameters of the biological system being modeled. Indeed, amino acid sequences and protein structures are on a different scale than the protein concentrations modeled in cell signaling. To overcome this challenge, we propose a ML pipeline that predicts the binding affinity parameters of cell signaling models using sequence and structure. We then compare these predicted parameters to the model parameters. ML models have been leveraged to similarly predict the parameters of models of cellular metabolism and steady-state signaling (8,9) (10). We note that the latter study on steady state signaling was tested on only one protein in the system, limiting the generalizability of the results. Excitingly, ours is the first such approach tested on dynamic signaling models. Furthermore, we aim to increase the generalizability of our results by presenting an expanded sample size.

Ultimately, we want to use our proposed framework to evaluate how much information is gained from adding sequence and structure measurements to our signaling model parameter inference. Despite interest in generalizing large datasets using ML models, there are a limited number of studies quantifying how much information is being gained from augmenting measurements traditionally used to calibrate systems biology models. For example, only recently has the limited applicability of the ML model used to predict parameters of cellular metabolism models outside of the ML model’s training data set been revealed (11). To fill this important gap, our proposed framework also comprises a generalizable Bayesian inference approach.

Bayesian inference is an established means to quantify the information gained from data. For example, one particularly relevant study used it to quantify the information gained from absolute measurements in the context of cell signaling models, which typically only leverage relative measurements for parameter inference (12). Importantly, Bayesian inference characterizes the posterior distribution, which is the distribution of plausible model parameters given a particular dataset. Inference begins by defining a prior distribution, the distribution of plausible model parameters given prior information. For example, the prior might formalize the fact that physical parameters can only be positive. The inference approach then proceeds by incorporating experimental datasets. To quantify the information gained from a particular dataset, one can compare the distribution of plausible parameter values before and after integrating data using statistical distance metrics. One common distance metric used to quantify the difference between distributions is Kullback Leibler (KL) divergence (13). The more the posterior distribution changes, the larger the KL divergence becomes, indicating that more information is gained from the data. KL divergence has been applied in experimental design and hypothesis generation (14–16). Here, we use it to quantify the information gained from incorporating experimental observations of sequence or structure into the parameter inference of cell signaling models.

Overall, we are proposing a novel, multiscale, probabilistic modeling framework that enables data integration across scales and quantifies the benefit of this integration (Figure 1). We use an ML pipeline to integrate multiscale data (amino acid sequence and protein structure) into the probabilistic, Bayesian parameter inference problem of ODE models of intracellular signaling. This probabilistic inference allows us to quantify the information gained from augmenting parameter inference with sequence and structure.

We test our approach on two, well characterized ODE models of cell signaling: Epidermal Growth Factor Receptor (EGFR) and G-Protein Coupled Receptor (GPCR) signaling. Not only are these models well-established; they are also biologically significant. In the case of EGFR signaling, this model represents dynamics upstream of the most well-studied pathway in cell signaling: MAPK (17). In the second case, modeling GPCR signaling is important as GPCR’s are considered a prime drug target due to their ubiquity and targetability. In fact, around a third of drugs approved by the U.S. Food and Drug Administration (FDA) target this protein (18).

To understand what information is gained from using sequence or structure in the parameter inference of GPCR and EGFR signaling, we compare the results of two parameter inference approaches. In what we will refer to as the “baseline approach”, only the published data reported for the EGFR or GPCR model is used to infer model parameters. In the “augmented approach”, the reported data is used together with data from PDB or UniProt. Data from PDB and UniProt is integrated using our multiscale probabilistic framework. Throughout the paper, we refer to these two datasets as *y_BASE_* and *y_AUG_*. We ultimately compare the predictions and posterior inferred using *y_AUG_* to the predictions and posterior inferred using *y_BASE_*. We perform this comparison using KL divergence, an information theoretic distance enabled by our probabilistic framework, to quantify the information gained from augmenting the baseline data. Ultimately, we answer the question of what information can be gained from protein structure and amino acid sequence in the context of dynamic models of cell signaling.

## 2 Approach and Results

### 2.1 Machine Learning K_D_ Predictor Pipeline

To leverage the large quantities of data on amino sequence and protein structure in the context of cell signaling models, we must translate the information in a sequence or structure to a scale that is meaningful for cell signaling — that is, on the scale of protein concentrations. Further, this translation approach should be easy to use to complement existing modeling approaches for cell signaling. We developed a ML pipeline for this task (Figure 2).

**Figure 2.**
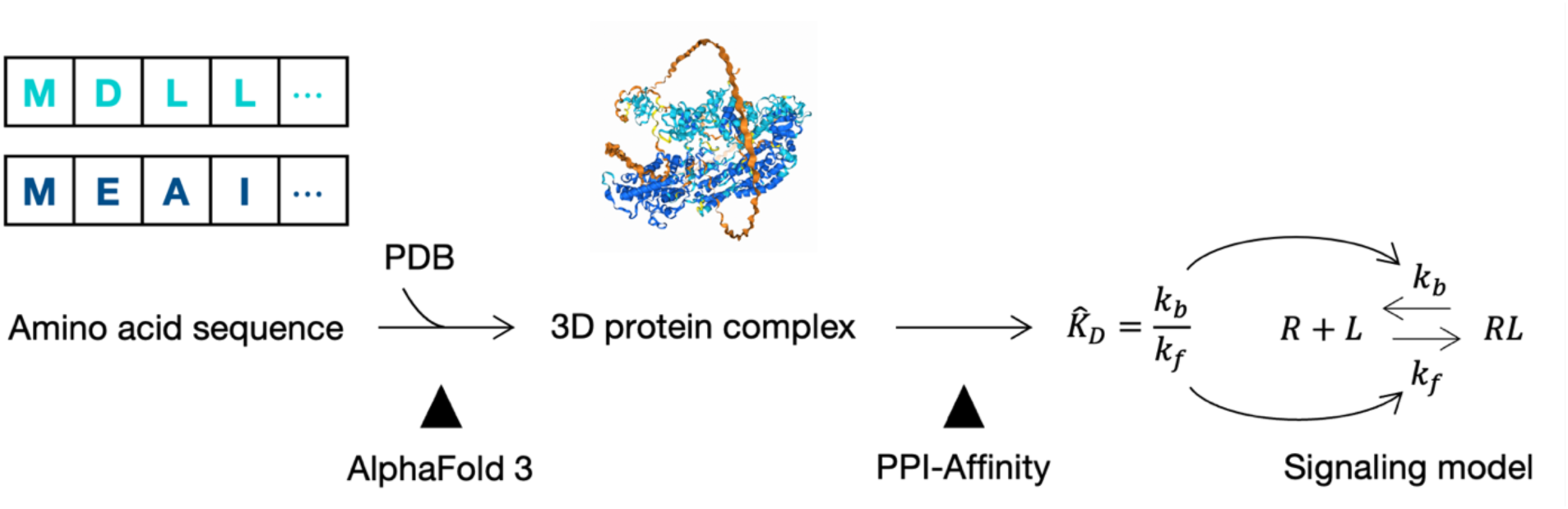
ML Pipeline to predict *K_D_*. *K_D_* is predicted using an SVM regression model, PPI-Affinity. PPI-Affinity needs the complex structure as an input. If the structure exists, it may be extracted from PDB. If it does not exist, the structure may instead be predicted using AlphaFold 3. AlphaFold 3 only needs amino acid sequences, which are typically available. The predicted *K_D_* is the same *K_D_* that characterizes reversible, bimolecular binding reactions comprising cell signaling models.

Upon reviewing the literature, we found that there exist several ML models that translate the information in an amino acid sequence and/or structure to a scale that is meaningful for cell signaling (19–22). Specifically, there are several ML models that predict protein binding affinity, *K_D_*, in units of concentration, from structure or sequence. We can then relate this predicted *K_D_* to the parameters of the ODE model, which are the unbinding (𝑘_𝑏_) and binding (𝑘_𝑓_) rates governing the reaction, by using the definition of *K_D_*: the ratio of the unbinding rate to the binding rate. These ML models are generally limited to bimolecular, reversible binding reactions. However, this is not a limitation in the context of cell signaling models, which are often comprised of such reactions. This is because even when the product of the reaction contains more than two proteins, the process usually involves a series of bimolecular reactions (23).

We found two ML models with convenient webservers for making *K_D_* predictions: Prodigy and PPI-Affinity (19,24). We moved forward with PPI-Affinity, a support vector machine (SVM) regression model, as it shows modest improvements on Prodigy’s performance. Further, it performs comparably to more complicated *K_D_* predictors, like those based on deep learning models (19). PPI-Affinity requires the structure of the bound protein complex as an input; however, only a fraction of experimentally available protein structures are complexes. Instead, most structures available are for monomer proteins. Thus, we sought an alternative input to PPI-Affinity to use in cases where the protein structure is not available.

Excitingly, AlphaFold (AF3), the latest iteration of the Nobel-prize winning, deep learning model AlphaFold, is designed to predict the structures of biomolecular interactions, including protein complexes (25). Although AF3 needs amino acid sequences of the binding partners to predict this structure, the amino acid sequence of a given protein of an intracellular signaling network is often known. AF3 is also hosted on a webserver, making predictions straightforward.

Altogether, the compatibility between signaling models and ML models, recent breakthroughs in predicting the structure of biomolecular interactions, and attention towards the useability of ML models enabled us to chain together existing ML models into one, user-friendly pipeline for incorporating amino acid sequence and protein structure into cell signaling models (Figure 2).

### 2.2 Test Case on Uninformative Prior

Before incorporating predictions from our ML pipeline into the parameter inference, we first tested its ability to accurately predict *K_D_* for the ten binding reactions present in the ODE test cases. For each of the ten binding reactions, we predicted *K_D_* using our proposed ML pipeline. We compared the accuracy of these predictions to an uninformed *K_D_* prediction. To generate this uninformed prediction, we sampled our Bayesian prior ten times. This prior is log-uniform, and has been used as an uninformative distribution for parameter inference in systems biology (12). Thus, it is well suited for representing an uninformed *K_D_* prediction. More details about this prior may be found in the Materials and Methods.

We calculate the absolute error of the ML pipeline predictions and prior samples with respect to the experimentally-derived *K_D_* values reported in the GPCR and EGFR publications (26,27). Since we are concerned with order-of-magnitude changes in *K_D_*, this error is calculated after a log10 transformation of the ML pipeline prediction and reported *K_D_*values. Figure 3a shows how the ML pipeline’s performance compares to the uninformed prediction’s performance. The ML pipeline’s predictions have a significantly lower absolute error than the uninformed prediction. One of these *K_D_* predictions was made with an experimental structure. Interestingly, it resulted in only the fifth smallest error.

**Figure 3.**
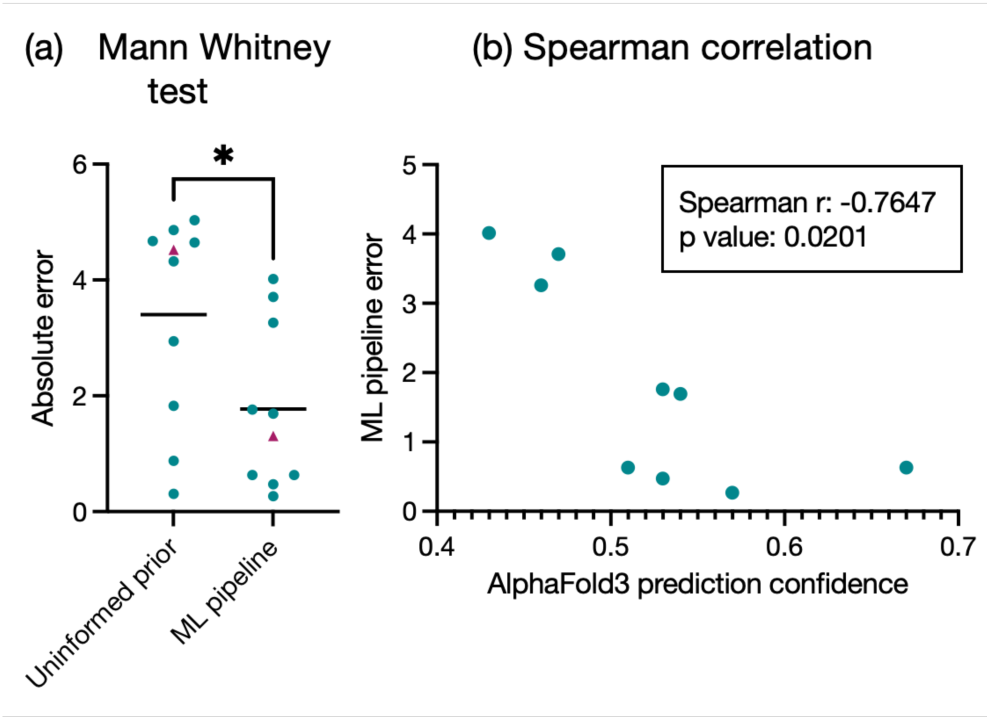
ML Pipeline performance. Cyan, EGFR reaction; pink, GPCR reaction. Circles, K_D_ predicted using predicted structure; triangles, K_D_ predicted using experimental structure. (a) Mann-Whitney test comparing absolute error of structure-informed K_D_ prediction with a random sample from a log-uniform distribution. n=10 binding reactions. *p-value< 0.05. Black horizontal line, sample mean. (b) Spearman’s rank correlation between error of K_D_ prediction and AlphaFold 3 confidence metric, ranking score. n=9 binding reactions that were predicted using a predicted structure, given there was no experimental structure.

We also tested whether the confidence metrics provided by the ML models that comprise the pipeline were indicative of the pipeline’s overall performance. To do this, we calculated the Spearman correlation between the error of the ML pipeline’s prediction and each confidence metric — the applicability domain (AD) for PPI-Affinity and the ranking score for AlphaFold 3. More details on these metrics can be found in the Materials and Methods. In the case of AF3, we test on only nine of the binding reactions. This is because for one of the reactions, there is an experimental structure. Thus, there was no need to use AlphaFold to predict the structure. We found that PPI-Affinity’s AD metric was not significantly associated with the overall pipeline’s performance (Supplementary Information Figure S1a). However, we did find that AlphaFold 3’s ranking score was significantly correlated with the pipeline’s performance (Figure 3b).

### 2.3 Test Case on EGFR and GPCR Signaling: Parameters

Next, we infer the Bayesian posterior distributions of the ODE model parameters using *y_BASE_* or *y_AUG_*. Details on this inference can be found in the Materials and Methods. We analyze the posterior samples after a log10 transformation, as we want to compare order-of-magnitude changes in the parameters. The quantiles of the resulting marginal posterior distributions of the unbinding and binding rates are shown in Figure 4a. After checking that both inferences recapitulate the training data (Supplementary Information Figure S3), we analyzed the resulting posterior distributions. To measure the information gained from using the augmented dataset compared to the baseline dataset, we compute the KL divergence from the posterior estimated with *y_BASE_* to the posterior estimated with *y_AUG_*. The resulting KL divergences are shown in Figure 4b. Interestingly, the most information is gained on the rate of protein unbinding. Figure 4c plots the mean absolute log error the parameter estimates with and without data augmentation. Error is calculated with respect to the experimentally derived reported values from the published papers. Excitingly, we see that the information in *y_AUG_*improves the parameter estimates, especially in the case of the unbinding rates, where the improvement is statistically significant.

**Figure 4.**
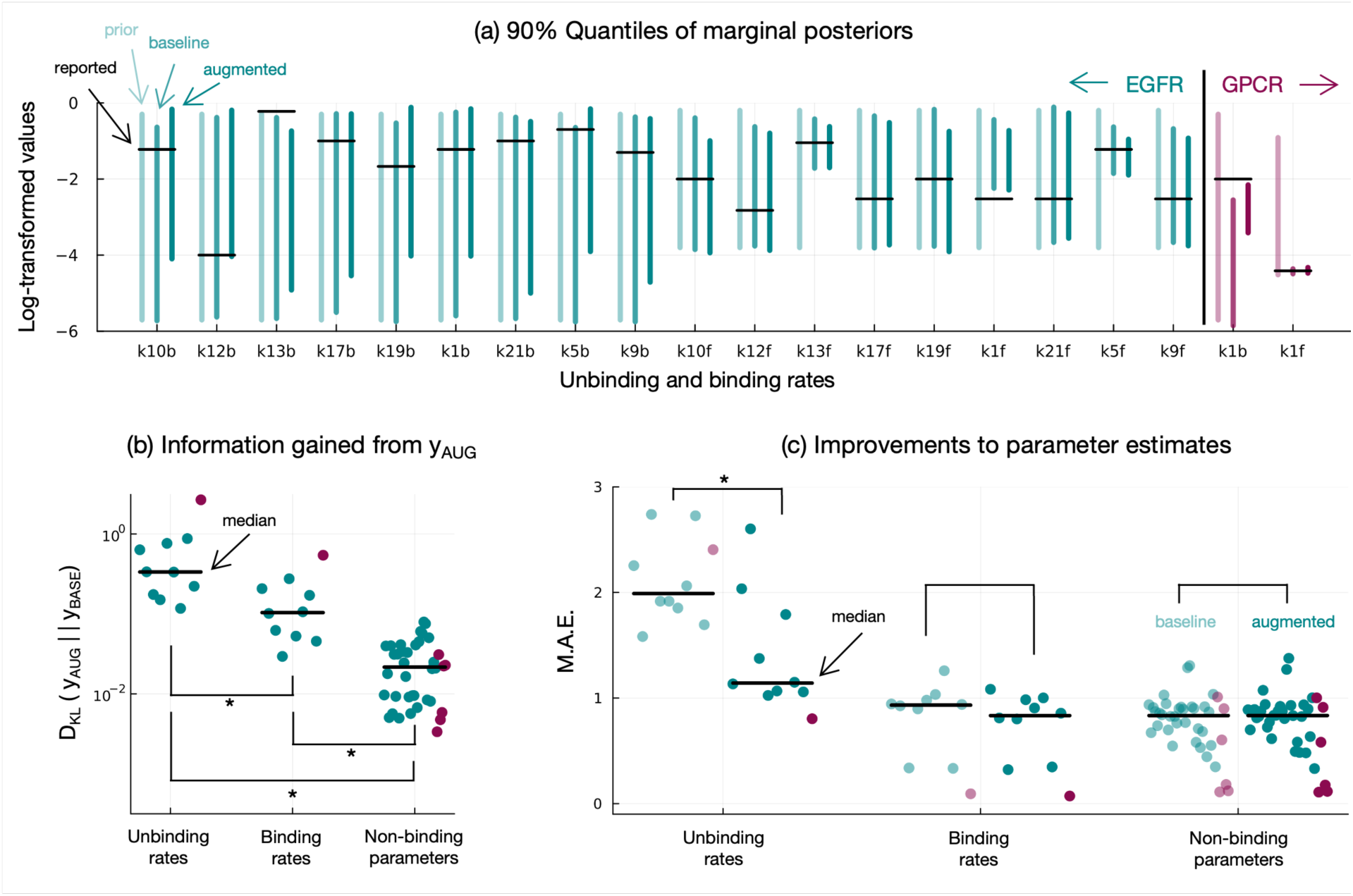
Parameter inference with and without data augmentation. Cyan, EGFR results; pink, GPCR results. All parameter samples were log-transformed prior to analysis. *p-value < 0.05 (a) 90% quantiles of marginal posterior distributions of binding parameters. Light cyan line, prior; medium cyan line, baseline posterior; dark cyan line, augmented posterior; black horizontal line, reported parameter value. (b) KL divergence, in bits, from baseline posterior to augmented posterior. Values are grouped by parameter function. (c) Mean absolute error (M.A.E.) of parameter samples. Mean taken with respect to each posterior distribution. Error calculated with respect to the values reported in the literature. Light cyan points, baseline; dark cyan points, augmented; black horizontal line, median M.A.E.

### 2.4 Test Case on EGFR and GPCR Signaling: Predictions

Figure 5 shows the difference in the medians and 90% quantiles of the test predictions generated by sampling the augmented posterior compared to the baseline posterior. The test/train split is detailed in the Materials and Methods. While we see some differences at middle timepoints of the EGFR test set (Figure 5a) and at certain doses of the GPCR test set (Figure 5b), overall the predictions are not qualitatively different. Further, in the case of GPCR, at no point does the difference between the medians of the two approaches become greater than the reported experimental error. In other words, the difference in the predictions between the two approaches falls within the error of the test set. In the case of EGFR, the difference between the medians does approach the level of the reported error. However, we note the unusually small scale of this reported error, which is +/- 1% change in phosphorylated SHC protein.

**Figure 5.**
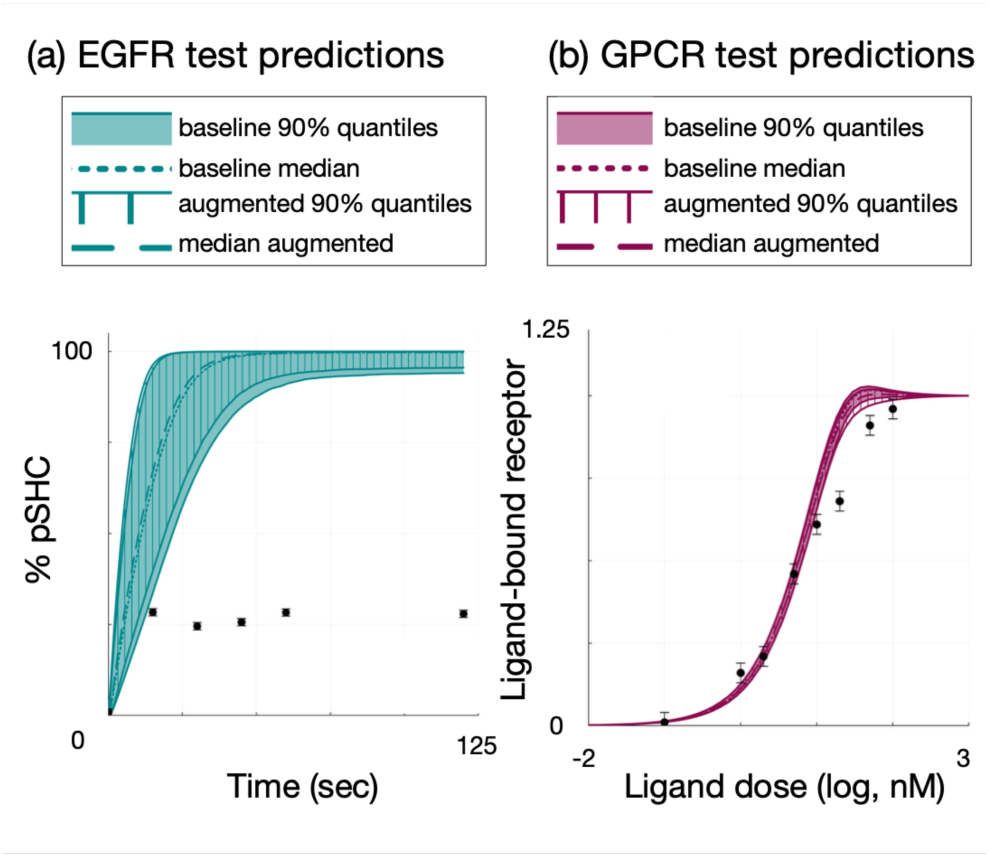
Performance on test set. (a) Predictions for EGFR test set, the percent of phosphorylated signaling protein SHC from 0-120 seconds. (b) Predictions for GPCR test set, the amount of ligand bound receptor 60 seconds post-stimulation, at different ligand doses, relative to 1000 nM of ligand. Cyan, EGFR; pink, GPCR. Shaded region, 90% quantiles of baseline approach; patterned region, 90% quantiles of augmented approach; dotted line, median prediction of augmented approach; dashed line, median prediction of augmented approach; black dots, experimental data with reported error.

When we investigate the medians and 90% quantiles for other outputs besides the test set, we observe larger differences in the predictions. For example, if we consider the timeseries of receptor-bound ligand, RL, of GPCR at times other than 60 seconds post-stimulation, we see greater differences between both the quantiles and medians (Figure 6a). Similarly, several species comprising EGFR signaling show larger differences in the predicted median and 90 percent quantiles. (Supplementary Information Figure S2).

**Figure 6.**
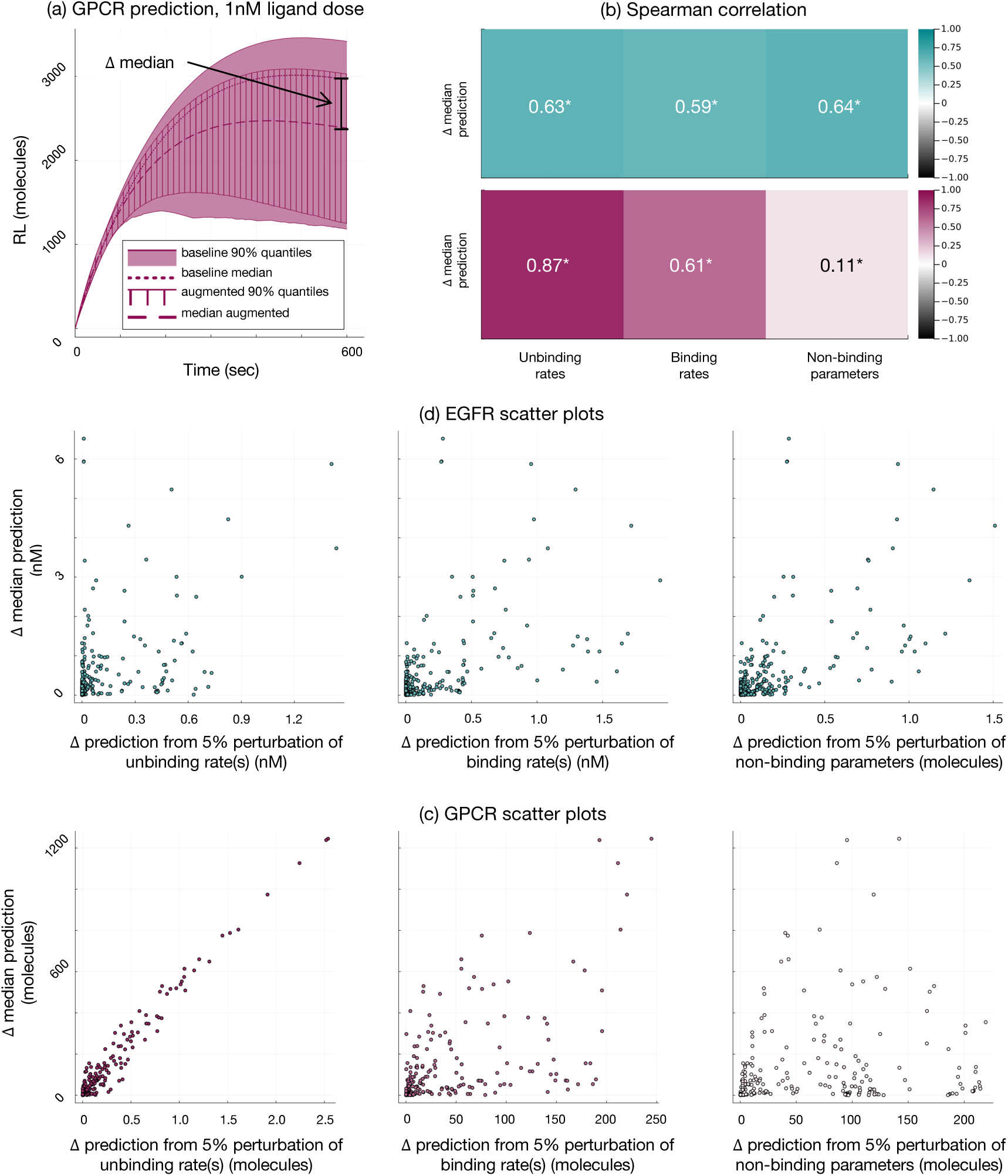
Local sensitivity analysis. (a) GPCR timeseries prediction of ligand-bound receptor, (RL), after 1 nM ligand stimulation. Units, molecules. Shaded region, 90% quantiles of baseline approach; patterned region, 90% quantiles of augmented approach; dotted line, median prediction of augmented approach; dashed line, median prediction of augmented approach. Difference between median prediction of baseline approach and median prediction of augmented approach, 𝛥 median, indicated with black arrow. (b) Spearman correlation between 𝛥 median and change in output due to 5% perturbation of either unbinding, binding, or non-binding parameters. Cyan, EGFR; pink, GPCR. *p-value< 0.05. n = 345 for EGFR; corresponding to 345 prediction outputs from 3 doses, 23 species, and 5 timepoints. n = 360 for GPCR; corresponding to 360 prediction outputs from 9 doses, 5 species, and 8 timepoints. Doses and timepoints match experimental observations. (c) Scatter plots illustrating relationship between perturbation and the 𝛥 median prediction for GPCR. (d) Scatter plots illustrating relationship between perturbation to a type of parameter and the 𝛥 median prediction for EGFR.

### 2.5 Local Sensitivity Analysis

To investigate what leads to changes in some, but not all, ODE model predictions, we conducted a local perturbation analysis of both the GPCR and EGFR outputs to model parameters. We perturbed the maximum likelihood parameter values of the baseline case by 5% and used the local sensitivity at the maximum likelihood to calculate the projected change in an output from this perturbation. We analyzed 360 outputs for GPCR and 345 outputs for EGFR. These outputs correspond to five model species at eight timepoints and nine doses and 23 model species at five timepoints and three doses, respectively for the two test cases. Doses and timepoints match experimental conditions. However, we exclude timepoint zero because the sensitivity of the initial concentrations to any parameter is zero. More details about the local sensitivity analysis may be found in the Materials and Methods section.

Next, we calculated the Spearman correlation between the change in the median prediction of an output and the sensitivity of that output to perturbations of either unbinding rates, binding rates, or non-binding parameters. The resulting Spearman coefficients for both EGFR and GPCR are shown in Figure 6b. Both test cases display a positive and significant correlation with the sensitivity to perturbation of unbinding rates. In the case of GPCR, the correlation decreases from unbinding to binding to non-binding parameters. This decreasing correlation was clear visually (Figure 6c). Interestingly, this is the case despite the sensitivity to perturbations being around an order of magnitude lower for unbinding rates than the other parameter types. In the case of EGFR, there is a significant positive correlation for all parameter types.

### 2.6 Investigation of Machine Learning Prediction Confidence

Given the significant correlation between the confidence of the protein structure prediction and absolute error of the predicted *K_D_*, (Figure 3b) we investigated whether weighing our data integration by the confidence of the predicted structure could improve our results. More details on how we implemented this confidence weight can be found in the Materials and Methods.

However, considering the confidence of the protein structure prediction did not significantly change most of the results of our parameter inference with and without data augmentation (Supplementary Information Figure S4). As in Figure 4, data augmentation provided the most information on unbinding rates. *y_AUG_* also consistently improved parameter estimates of unbinding rates compared to *y_BASE_*.

Predictions of the test data were similarly unchanged qualitatively (Supplementary Information Figure S5). As in Figure 5, the predictions with and without data augmentation were similar. One quantitative difference did emerge. When structure confidence was considered, the difference between median predictions for the EGFR test data was no longer greater than the reported error.

### 2.7 Robustness of Results to Changes in Prior Distribution

The prior distribution for each binding parameter is representative of a non-informative characterization of our knowledge of binding and unbinding rates. However, we also wanted to explore how the results might change given a more informative or less informative prior. To do so, we increased or decreased the upper and lower bounds of the prior distributions of all parameters by one order of magnitude. The results were qualitatively consistent across these changes in the prior (Supplementary Information Figures S6, S7, S8, and S9). Further, they were mostly quantitatively consistent as well; only a few quantitative differences emerged. In the case of the more informative prior, the difference between median predictions for the EGFR test data was no longer greater than the reported error (Supplementary Information Figure S7a). In the case of the less informative prior, the information gained from data augmentation was no longer significantly different between the unbinding and binding rates (Supplementary Information Figure S8b). This was because more information was gained for binding rates. Interestingly, this did not result in a corresponding significant change in the mean absolute error for binding rates.

## 3 Discussion

Computational models of biological systems generate insights beyond what experiments alone can offer. However, they also present their own challenges. Our work here explores the potential of augmenting the conventional measurements used for parameter inference in models of intracellular signaling. Ultimately, our goal was to reduce uncertainty in parameters for better model determination. To do this, we proposed a multiscale, probabilistic modeling framework. We first develop a ML pipeline for incorporating the new, multiscale measurements into parameter inference. Then, we conduct a thorough quantification of the information gained from these new measurements using a Bayesian framework.

We use amino acid sequence and protein structure measurements from UniProt and PDB, respectively, to augment the published datasets of two test cases: EGFR and GPCR signaling. The published datasets include partial, relative measurements of protein concentration and protein co-localization over time. We chose the EGFR and GPCR models as our test cases because both are well-established and biologically significant. Taken together, these models offer a comprehensive test of our approach. For example, EGFR provides a test of a moderately sized model, while GPCR provides a test of a minimal model. In addition, our ML pipeline can make *K_D_* predictions using sequence or structure. EGFR provides a test of predictions using sequence, while GPCR provides a test of predictions using structure.

Predictions from the GPCR and EGFR models occur on the scale of protein concentrations. To make measurements of sequence and structure applicable to the parameter estimation of these models, we needed to convert the measurements to the same scale as the GPCR and EGFR models. Excitingly, we were able to integrate two, user-friendly ML models, PPI-Affinity and AlphaFold 3, into one pipeline to do this (19,25). This pipeline uses sequence or structure to predict the binding affinity, *K_D_*, of a given bimolecular, reversible reaction in a cell signaling model. We find that this ML pipeline outperforms an uninformative prior when predicting the *K_D_*. Thus, it is appropriate to use when no experimental measurements exist for *K_D_*. Importantly, this pipeline is generalizable to other dynamic signaling models. It is limited only by access to amino acid sequence and comprises two, user-friendly web servers.

Interestingly, we found that our pipeline’s performance did not significantly benefit from using an experimental protein complex structure, rather than predicted complex structure, when predicting *K_D_*. The binding affinity between GPCR and its ligand was predicted using an experimental structure. However, the error of this prediction was not smaller than all the other *K_D_* prediction errors. PPI-Affinity was trained using complex structure and *K_D_* pairs from the PDBbind database. Although the GPCR structure was within the applicability domain of PPI-Affinity, it seems that the ligand-bound GPCR complex was not included in this training set. This could explain the lack of improvement we see. We also note the small sample size (n=1) of experimental structures we tested, which limits the certainty of this result.

We incorporate the *K_D_*’s that we predict using the ML pipeline into the likelihood that we use in our signaling model’s parameter inference. We combined this likelihood of the predicted *K_D_* and the likelihood of the data reported in the original publications to form an augmented dataset, y_AUG_. This augmented dataset is ultimately informed by the experimental data on sequence or structure that we use to predict *K_D_*. We then compared the posteriors and predictions inferred with y_AUG_ to the posteriors and predictions inferred using only the original data, y_BASE_, our baseline dataset.

Overall, augmenting our data with data on sequence or structure provides more information on rates of protein unbinding compared to using only the original data. This was consistent across both the EGFR test case and GPCR test case. Further, the information gained from sequence or structure tends to push the unbinding rate towards the rates reported in the original publications. These reported rates were all taken from experiments, rather than fit to data (26,27). Our results offer an understanding of when our approach is useful: in the case of underdetermined rates of protein unbinding. We have confidence in this recommendation of utility, since the results were consistent even with a more informative or less informative prior distribution. In practice, it is helpful to use our approach to constrain unbinding rates, as these rates tend to vary over a larger range than binding rates (23).

When we investigate how predictions change due to the data augmentation, the results are less clear. We first find that augmenting the baseline data set does not fundamentally change the predictions of our test data for either test case. However, we do see more substantial changes for other outputs besides the test data. We thus investigated when predictions inferred with *y_BASE_* will differ from predictions inferred with *y_AUG_*. As our approach changes unbinding parameter rates, intuitively, one would then expect to see differences in predictions for outputs influenced by unbinding rates. Thus, we investigated the relationship between the local sensitivity of an output to unbinding rates, binding rates, and non-binding parameters. When we analyzed the GPCR model, there emerged a striking, significant, positive and linear correlation between changes in a prediction and the local sensitivity of that prediction to perturbations in the unbinding rate. A significant positive correlation also emerged with the EGFR model. However, this relationship was similar across unbinding rates, binding rates, and nonbinding parameters. The different results from the two test cases may be a function of the complexity of the models. While GPCR is a minimal model comprised of eight parameters, EGFR is a more moderately sized model comprised of 50 parameters. It is thus plausible that the relationship between sensitivity and predictions is impacted by higher-order interactions between parameters in the case of EGFR (28). These interactions would not be captured by the local sensitivity analysis we conduct here.

There was a significant negative correlation between AlphaFold 3’s confidence metric and the error of the *K_D_*prediction. Based on this, we investigated weighing the likelihood of *K_D_* by AlphaFold’s confidence. Interestingly, incorporating confidence in this way did not significantly improve or change our results. This could be explained by the fact that we were conservative when balancing the influence of data augmentation against the influence of the baseline data. For example, weighing by confidence, which was scaled from [0,1] decreased the influence of predicted *K_D_*’s. An interesting future direction could be investigating other probabilistic frameworks for incorporating AlphaFold’s confidence into parameter inference.

We acknowledge some limitations of our study. We test our ML pipeline on only the 10 binding reactions reported in our test cases. We could test our pipeline with more reactions from databases of paired structures and affinities, like PDBbind (29). We chose to focus only on binding affinities reported in the context of ODE models as these affinities are known to recapitulate intracellular protein dynamics. Future work could investigate how our ML pipeline could be even more specialized to binding reactions occurring inside the cell. For example, future work could take in to account the crowded intra-cellular environment (30). With future improvements to the ML pipeline in mind, we developed a Bayesian framework for incorporating 𝐾_𝐷_ that can evolve with our pipeline. For example, we use the error reported by PPI-Affinity to weigh the predicted *K_D_* likelihood against the likelihood of the conventional experimental measurements. This weight can be adjusted as that error improves.

There are also ML models that use amino acid sequence to predict binding affinity. In theory, we could use one of these models instead of our pipeline. However, in practice, we found that there were no such models hosted on convenient webservers. Even more limiting was the prohibitively large RAM these models needed to make a prediction for the relatively large signaling protein complexes we are interested in here (20). We instead focus on more user-friendly models, both in terms of RAM and implementation.

In conclusion, we have developed a multiscale, probabilistic framework to show that it is viable to use measurements of amino acid sequence and protein structure to augment parameter inference in the context of dynamic signaling models. Experimental data on protein structure and amino acid sequence consistently adds information on the unbinding rate parameters of these signaling models. This improves parameter estimation and thus model determination. In the future, our framework has the potential to synergize with breakthroughs in ML and data collection. As ML models better predict binding affinities, our framework can better constrain signaling models. As databases of protein structure and amino acid sequence grow, our framework will generalize to more binding reactions.

## 4 Materials and Methods

### 4.1 ODE models of intracellular signaling

We base our investigation on two ODE models: a 50 parameter, 23 species model of EGFR signaling and an eight parameter, seven species model of GPCR signaling (26,27). In the case of EGFR signaling, a combination of mass action and Michaelis Menten kinetics is used to describe cell signaling events such as protein binding and phosphorylation. In the case of GPCR signaling, mass action kinetics are used to describe cell signaling events such as protein binding, activation, and degradation. Binding reactions are of particular interest to our study: in the case of EGFR, 18 parameters characterize nine binding reactions, while in the case of GPCR, two parameters characterize one binding reaction.

The EGFR model simulates signaling up to 120 seconds, while the GPCR model simulates signaling up to 600 seconds. We use the reported initial species pools to initialize the model simulations. We simulate both models with the Julia package *DifferentialEquations.jl* (31).

Because our model parameter values can vary across a wide range, the solution may be stiff. Thus, we want to choose an ODE solver that can handle stiffness. In addition, we need a solver that performs well given a higher-than-default accuracy requirement. This is due to the nature of our state variable: protein concentration. The accuracy of a particular problem is specified by the absolute and relative tolerances. The relative tolerance (rtol) is set based on the number of significant digits required for the model output, plus 1 (32,33). We note that the upper bound on the number of significant digits is between 15 and 17, as this is the maximum number of significant digits stored by a Float64 type variable, which we use here. In our case, we deem 5 significant digits sufficient for our problem and set the rtol to 10^-(5+1)^. The absolute tolerance (atol) is set based on the absolute value of the model output that may be deemed insignificant (32,33). In our case, we deem a protein concentration of 10^-5^ nM, or less than 1 protein/cell, as insignificant. A rtol of 10^-6^ and atol of 10^-5^ are more as stringent as the default tolerances for most scientific computation, 10^-3^ and 10^-6^, respectively (32,34). Based on our requirements, we choose to use *DifferentialEquation.jl*’s QNDF solver, which is equivalent to Matlab’s ode15s solver (34). It is a stiff solver that can handle medium tolerances; thus, it meets our problem’s requirements (32,34,35).

### 4.2 Data Augmentation

Given an ODE model, 𝑀(·), and experimental data, 𝑦, our objective is to infer the model parameters, 𝜃. In this work, we propose augmenting the published data for each system, *y_BASE_*, with predicted *K_D_* values, 𝐾̂_𝐷_. Equation (1) shows how the augmented dataset, *y_AUG_* incorporates additional data into *y_BASE_*. We predict *K_D_* using experimental observations of amino acid sequences from UniProt, 𝑦_𝐹𝐴𝑆𝑇𝐴_, or protein structures from PDB, 𝑦_𝑃𝐷𝐵_. *y_BASE_* , 𝑦_𝐹𝐴𝑆𝑇𝐴_ and 𝑦_𝑃𝐷𝐵_ are defined in detail in the subsequent sections.

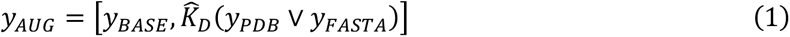

Throughout the paper, we refer to 𝐾̂_𝐷_(𝑦_𝑃𝐷𝐵_, 𝑦_𝐹𝐴𝑆𝑇𝐴_) as simply predicted *K_D_*, for conciseness and clarity.

### 4.3 Experimental measurements of EGFR and GPCR signaling

Both ODE models report experimental measurements of the system of interest. That is, measurements of in vitro protein dynamics, which correspond to the predictions of the ODE models (Figure 7). Note that we are not pooling data across systems, but, for simplicity, we will refer to each data set as *y_BASE_*. In the context of the EGFR model, *y_BASE_* are measurements of EGFR signaling dynamics (Figure 7a) and in the context of the GPCR model, *y_BASE_* are measurements of GPCR signaling dynamics (Figure 7b). We extracted the data and errors with Plot Digitizer (36) .

**Figure 7.**
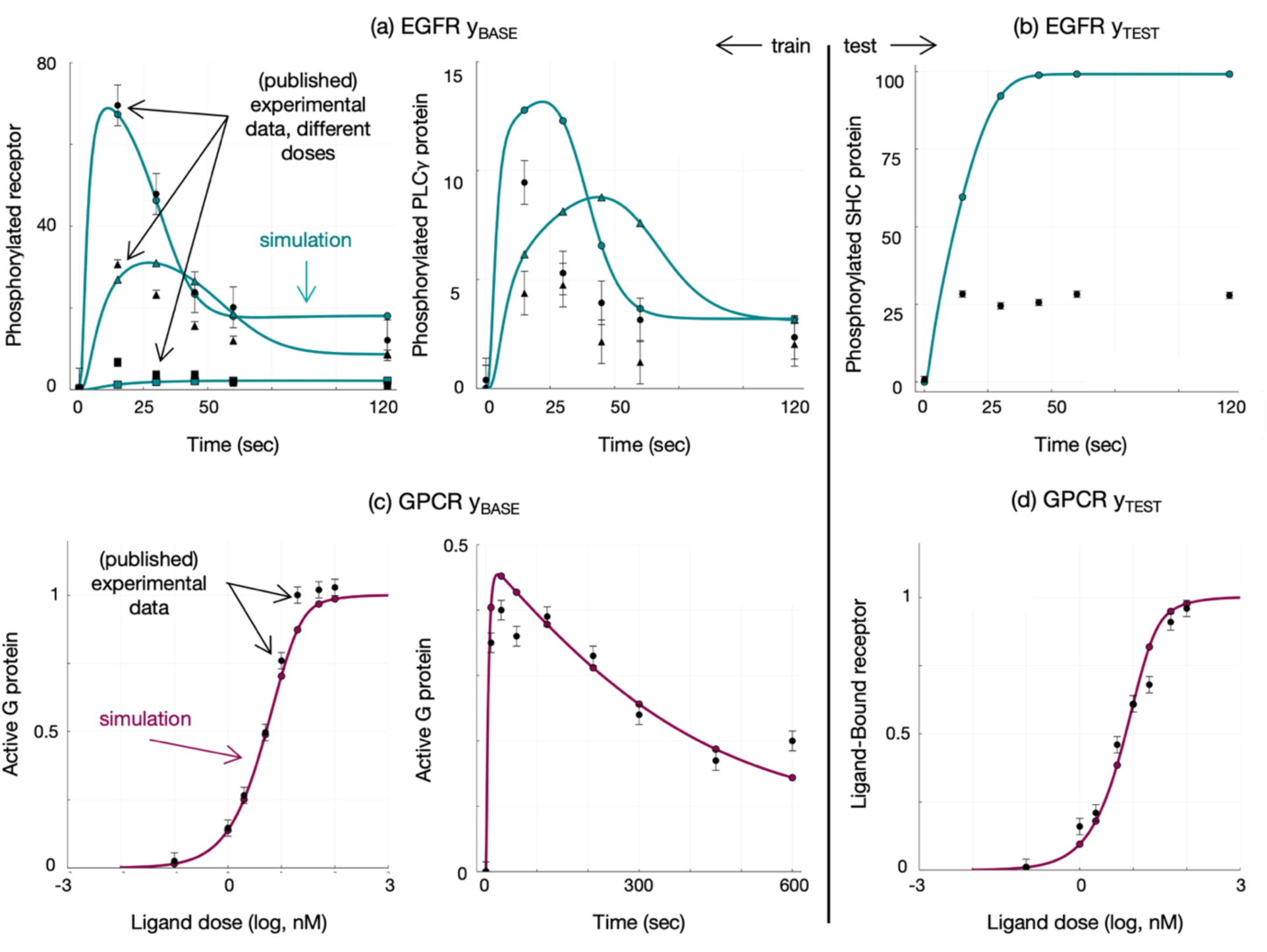
Baseline data set (*y_BASE_*). (a) Training data for EGFR parameter inference. (b) Test data for EGFR parameter inference. (c) Training data for GPCR parameter inference. (d) Test data for GPCR parameter inference. Cyan or pink lines, simulations using reported parameter values for EGFR and GPCR, respectively; black markers, reported experimental data points for a particular dose; black error bars, reported standard error.

Eight outputs of EGFR are observed, and we show three of these outputs as a representative sample in Figure 7a. These include the percent of phosphorylated receptor at various ligand dosages as well as the percent of phosphorylation of two intracellular proteins at various ligand dosages. Further observations include the percent of an intracellular protein bound with another intracellular protein and the receptor. The original work does not fit parameters to data, so we must decide our test/train data split. We choose to reserve measurements of one of the intracellular proteins, SHC, as our test data. We choose SHC due to its clinical relevance, as this signaling species has been shown as an important marker in various cancers (37). Furthermore, the published model predictions do not quantitatively match the experimental measurements of SHC (37). Ideally, incorporating data on protein structure will improve our predictions for this experimentally relevant target. Thus, we use the data for SHC to test our model predictions.

Three outputs of GPCR signaling are observed, all of which are shown in Figure 7b. These include levels of active G-protein at various ligand dosages as well as levels of the ligand-bound receptor at various ligand dosages. As in the original work, we use the two G-protein data sets as training data. We reserve data on the bound receptor for testing.

Both models also report binding parameter values derived from experiments or physical bounds, rather than fit to the data. We use these reported values as a test of the binding parameter values themselves.

### 4.4 Experimental measurements from UniProt and PDB Databases

We augment *y_BASE_* with amino acid sequences or structures of the proteins that are involved in binding for each system. Given two proteins that are binding, we first searched for any structures of the bound complex in PDB. One such structure exists, describing GPCR bound to alpha pheromone, PDB structure 7QBI (38,39). This structure corresponds to the single binding reaction in the GPCR model and is formatted as a PDB file. This measurement comprises 𝑦_𝑃𝐷𝐵_.

If no such structure was found, as was the case for the nine EGFR binding reactions, we extracted the canonical amino acid sequences of the individual proteins, as FASTA files, from UniProt. They are extracted UniProt sequences P01133 (EGF), P29353 (SHC1), P19174 (PLCG1), Q07889 (SOS1), P62995 (GRB2), and P00533 (EGFR). These measurements comprise 𝑦_𝐹𝐴𝑆𝑇𝐴_.

### 4.5 ML pipeline to predict 𝑲_𝑫_

Figure 2 provides an overview of the ML pipeline used to predict *K_D_* using amino acid sequence (𝑦_𝐹𝐴𝑆𝑇𝐴_) or protein structure (𝑦_𝑃𝐷𝐵_). Given either the sequences of the binding proteins or the structure of the bound complex, our objective is to predict the binding affinity, *K_D_*, in units of concentration, for a specific bimolecular binding reaction. We can then relate this predicted *K_D_* to the parameters of the ODE model, which are the unbinding (𝑘_𝑏_) and binding (𝑘_𝑓_) rates governing this reaction, by using the definition of *K_D_*: the ratio of the unbinding rate to the binding rate.

If the bound complex structure is available, we can use a support vector machine (SVM) regression model, PPI-Affinity, to predict Δ𝐺, the free energy change in kcal/mol (19). PPI-Affinity uses a feature vector derived from the complex structure to make its prediction of Δ𝐺. We then convert Δ𝐺 to *K_D_*, given a gas constant and absolute temperature (40,41). If the bound complex is not available, we use a deep-learning model, AlphaFold 3, to predict the complex structure given the amino acid sequences of the individual proteins. If the complex is comprised of identical subunits, we extract the smallest subunit (21). We then use this predicted structure as in input to PPI-Affinity to predict *K_D_*.

Both PPI-Affinity and AlphaFold 3 provide confidence metrics for their predictions. AlphaFold provides a ‘ranking score’ for each predicted structure. This score ranges from [-100, 1.5] with - 100 representing low confidence. The ranking score is a function of various features of the protein structure, such as the fraction of the predicted structure that is disordered and the interface predicted template modeling (ipTM) score (25). Both the ranking score and ipTM are significantly correlated with the performance of the ML pipeline (Figure 3b and Supplementary Information Figure S1b). Conveniently, ipTM is scaled between [0,1], with zero representing low confidence. We take advantage of this property when considering AlphaFold 3 confidence in our Bayesian framework, detailed next. In the case of PPI-Affinity, it is noted whether each *K_D_*prediction is within the applicability domain of the training dataset (19). We found that the accuracy of predictions was more dependent on ranking score than applicability domain.

### 4.6 Bayesian inference

The objective of Bayesian inference is to find the posterior distribution, 𝑝(𝜃|𝑦) , which is defined using Bayes rule:

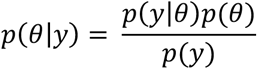

While we often know the likelihood of the data given model parameters, 𝑝(𝑦|𝜃), and the prior probability of parameters, 𝑝(𝜃) we usually do not know the probability of the data 𝑝(𝑦). Thus, we must approximate this distribution knowing 𝑝(𝜃|𝑦) up to a proportionality constant. While there are various distribution approximation methods for this problem, the most powerful are Markov-Chain Monte Carlo (MCMC) approximations, because our initial guess of 𝑝(𝜃|𝑦) does not need to be particularly close to the reality. These algorithms generate correlated samples of the posterior, which we may use to simulate our model.

To generate these samples for our ODE models, we use a Julia implementation of the affine invariant ensemble MCMC sampler from the package *AffineInvariantMCMC.jl* (42). Like all MCMC algorithms, the affine-invariant ensemble sampler algorithm constructs a Markov chain that is used to sample the posterior distribution. To better navigate the parameter space, this algorithm initializes an ensemble of samplers that propose the next sample based on the location of another sampler in the ensemble. Because all our parameter values are restricted to positive values and are often described on a logscale rather than linear scale, we conduct sampling in a log-transformed parameter space (43,44). Further discussion of the convergence diagnostics of the samples to the posterior may be found in the Supplementary Information.

### 4.7 Prior

The prior encodes our assumptions, ideally derived from empirical knowledge, about parameters. Here, we choose to define independent, log-uniform priors. If the parameter is a binding rate, we limit the distribution to [10^-4^ nM^-1^s^-1^, 100 nM^-1^s^-1^]; for an unbinding rate, we limit the distribution to [10^-6^ s^-1^, 100 s^-1^]. This is in line with prior distributions for binding rates in the literature and database values (12,45). For non-binding parameters, we limit the distributions to +/- two orders of magnitude above and below the reported parameter value.

### 4.8 Likelihood

Recall that our augmented data, 𝑦, is comprised of two categories, *y_BASE_* and predicted *K_D_*, 𝐾̂_𝐷_. Both *y_BASE_* and 𝐾̂_𝐷_ represent vectors of data points. For example, in the case of EGFR, *y_BASE_* is a vector of 42 points, corresponding to seven different outputs observed at six timepoints each, while 𝐾̂_𝐷_ is a vector of nine predicted affinities for the nine binding reactions of the model.

We assume that each data point is independent from the rest. Next, we assume that a data point in *y_BASE_*, 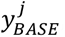, is normally-distributed, given the ODE model prediction and experimental error for data point j. For example, if 𝑗 indicates the amount of active G-protein 300 seconds after stimulation, then we use the model’s prediction for active G-protein 300 seconds after stimulation as the mean of the likelihood distribution and the reported error of this measurement as the deviation of the likelihood distribution.

Finally, we assume that a data point in 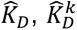 is log-normally distributed, given the ratio of the unbinding and binding reaction rates for that data point and standard error of the SVM regression model, PPI-Affinity. For example, if 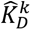 is the predicted binding affinity of a receptor binding a ligand, then we will use the rate of receptor ligand unbinding divided by rate of receptor ligand binding as the mean of the likelihood distribution and the standard error as the deviation. This standard error is on the scale of one order of magnitude change in 𝐾̂_𝐷_.

For parts of the investigation, we augment this likelihood term to account for AlphaFold3’s confidence. Specifically, we weigh the log-likelihood of a given 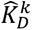 by AlphaFold 3’s interface predicted template modeling (ipTM) score, which represents the confidence of 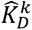 scaled from zero to one. This likelihood formulation is known as a power prior. Power priors consist of raising the likelihood, or a term in the likelihood, to a discounting factor. This controls the amount of information taken from certain data (46,47). When the log likelihood is evaluated, the product of likelihoods becomes a sum, and the discount factor(s) become weight(s) on the terms of this sum. Further details on our likelihood distribution are included in the Supplementary Information.

### 4.9 Log Transformation of Parameters for Analysis

We note that analyses of the model parameters, and therefore posteriors, are done after a log10-transformation of the parameter samples. The reason for this is threefold. First, log-transformation tends to bring biochemical parameters like *K_D_* towards a Normal distribution, which is desirable for most statistical analyses (43). Second, by comparing order-of-magnitude changes in parameters, we can compare parameters of different scales. Finally, we are concerned with order-of-magnitude changes in *K_D_*, as it has been shown that changes less than this do not significantly influence model predictions (48).

### 4.10 Quantifying information gained using KL divergence

We use KL divergence between marginal posterior distributions to quantify the information gained from augmenting our data with experimental measurements of amino acid sequence and protein structure. Specifically, we calculate the KL divergence from the marginal probability of parameter 𝜃_𝑖_ given only the baseline data, 𝑞(𝜃_𝑖_|*y_BASE_*), to the marginal posteriors estimated with the augmented data, 𝑝(𝜃_𝑖_|*y_AUG_*):

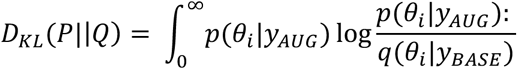

Given posterior samples, we use a univariate kernel density estimate and Monte Carlo integration to approximate a divergence for each marginal posterior (15).

### 4.11 Sensitivity Analysis and Perturbation Analysis

We compute the local sensitivity of a model prediction with respect to model parameters using Julia’s *ForwardDiff.jl* packages (49–51). Given the local nature of this analysis, parameters are not log-transformed. *ForwardDiff.jl* approximates the derivative of an output with respect to a model parameter using forward mode automatic differentiation. We use forward mode automatic differentiation as it is recommended for ODE’s with less than 100 parameters (49). We take the local sensitivity with respect to the posterior parameter set that maximizes the likelihood of the baseline training data.

To calculate the projected change in an output due to a 5% change in a particular parameter, we multiple the local sensitivity by 5% of the maximum likelihood parameters. We take the absolute value of the resulting perturbation. We then need to down select one perturbation value for each type of parameter: unbinding, binding, or non-binding. In the case of GPCR, this is trivial for unbinding and binding rates, as there is only one unbinding parameter and one binding parameter. For the other cases, GPCR non-binding parameters and all EGFR parameters, we take the maximum resulting perturbation.

### 4.12 Statistical Analyses

All statistical analyses, Mann Whitney U test and Spearman rank correlation, were carried out using Prism 10.4.0 or a combination of Julia’s *HypothesisTests.jl* package and *StatsBase.jl* packages.

### 4.13 Code Availability

Code is available at: https://github.com/FinleyLabUSC/structure_informed_cell_signaling.

## Acknowledgements

The authors thank members of the Finley research group for critical comments and suggestions. This work was partially supported by the National Cancer Institute of the National Institutes of Health grant 1U01CA27580 and National Science Foundation award 2410753.

