## Supplementary Information for "Multiscale Probabilistic Modeling: A Bayesian Approach to Augment Mechanistic Models of Cell Signaling with Machine-Learning Predictions of Binding Affinity"

### 1. Supplementary Methods – Details on Bayesian Inference

We use 1000 walkers to sample the 8-parameter GPCR model and 50-parameter EGFR model. This was a conservative number, chosen to exceed the minimum number of walkers for sampling, which is 2*number of parameters (1). These walkers are dependent, thus traditional metrics comparing variance within and between chains are not applicable (2).

#### 1.1 Convergence Diagnostics

We use the effective sample size, ESS, to monitor convergence to the posterior. ESS is a function of the autocorrelation of samples generated by the MCMC algorithm. Monitoring autocorrelation is the recommended metric for the affine invariant sampler, thus, we use ESS here (2). We conclude convergence once there are 100 effective samples per ensemble chain (3).

#### 1.2 $K_D$ Likelihood Standard Deviation

We use the M.A.E. reported by PPI Affinity as the standard deviation for the augmented likelihood term. PPI Affinity reports the mean absolute errors, M.A.E., in kcal/mol, for two test sets. One test set is comprised of protein-peptide binding reactions. The M.A.E. for this set is 1.1 kcal/mol. Another test set is comprised of protein-protein binding reactions. The M.A.E. for this set is 1.8 kcal/mol (4). We use the first M.A.E. as the standard deviation for the likelihood of $K_D$s characterizing protein-peptide binding reactions and the second M.A.E. as the standard deviation for the likelihood of $K_D$s characterizing protein-protein binding reactions.

To use the reported M.A.E., we need to convert from a change in energy in kcal/mol to a change in the binding affinity. A change in energy of about 1.35 kcal/mol results in an order of magnitude, or 10x, change in the binding affinity at room temperature (5). However, we are concerned with changes at body temperature—in that case, a change in energy of about 1.41

kcal/mol results in an order of magnitude change in $K_D$. Using this relationship, we calculate the order of magnitude change in $K_D$ given the changes in energy reported by PPI Affinity:

$$\frac{1}{1.41} = \frac{\sigma_{pep-prot}}{1.1} => \sigma_{pep-prot} = 0.8$$

$$\frac{1}{1.41} = \frac{\sigma_{prot-prot}}{1.8} => \sigma_{prot-prot} = 1.3$$

The augmented likelihood is on a log 10 scale. Thus, there are no further conversions needed to use these values in our likelihood. Overall, this means that we penalize the augmented likelihood term on the scale of one order of magnitude change in $K_D$—slightly less in the case of a peptide-protein interaction and slightly more in the case of a protein-protein interaction.

### 2. Supplementary Figures

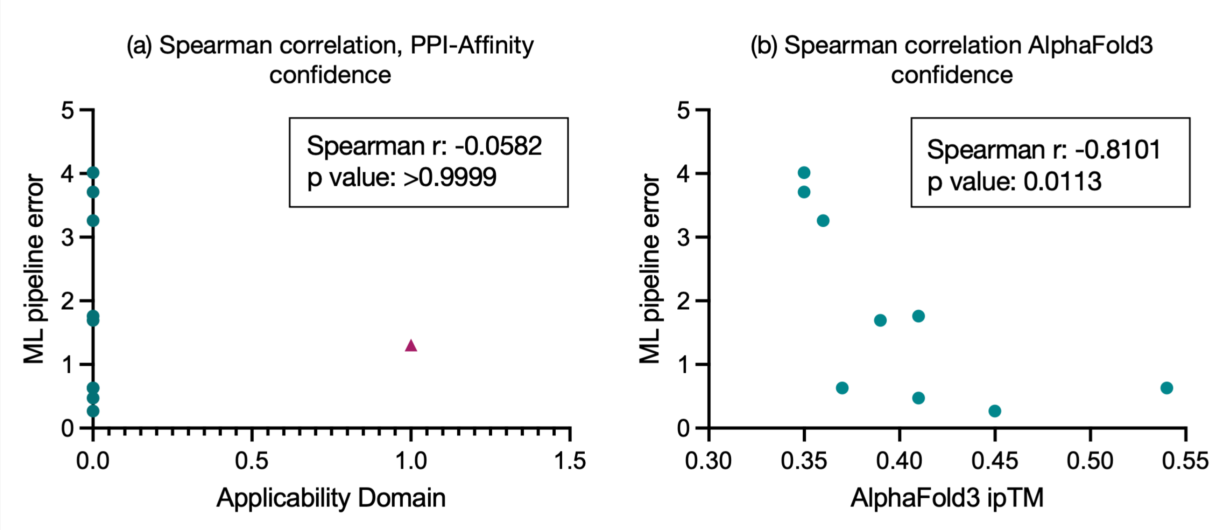


**Figure S1. ML Pipeline extended performance.** Cyan, EGFR reaction; pink, GPCR reaction. Circle, predicted structure, triangle, experimental structure. (a) Spearman's rank correlation between error of $K_D$ prediction and PPI-Affinity confidence metric, Applicability Domain. Applicability Domain is a binary metric, with 1 indicating confidence. n = 10 binding reactions. (b) Spearman's rank correlation between error of $K_D$ prediction and AlphaFold 3 confidence metric, ipTM. ipTM bounded from [0,1]. n=9 binding reactions for which we used predicted structure, given there was no experimental structure.

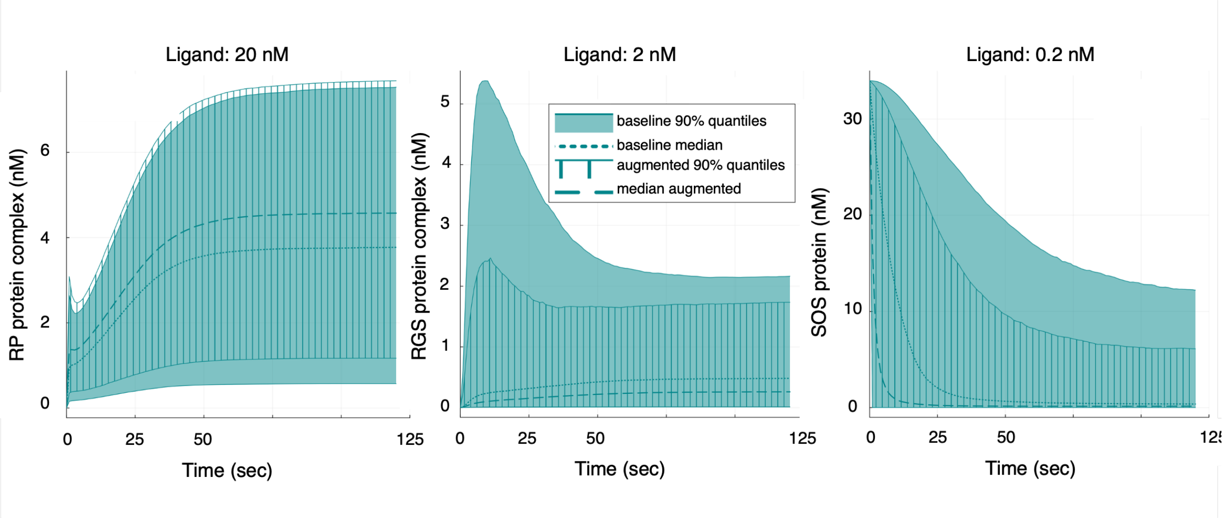


**Figure S2. EGFR timeseries predictions across time, ligand dose, and species.** RP, phosphorylated, ligand-bound receptor; RGS, phosphorylated, ligand-bound receptor bound to intracellular proteins GRB2 (G) and SOS (S). Prediction in nanomolar concentration. Shaded region, 90% quantiles of baseline approach; patterned region, 90% quantiles of augmented approach; dotted line, median prediction of augmented approach; dashed line, median prediction of augmented approach.

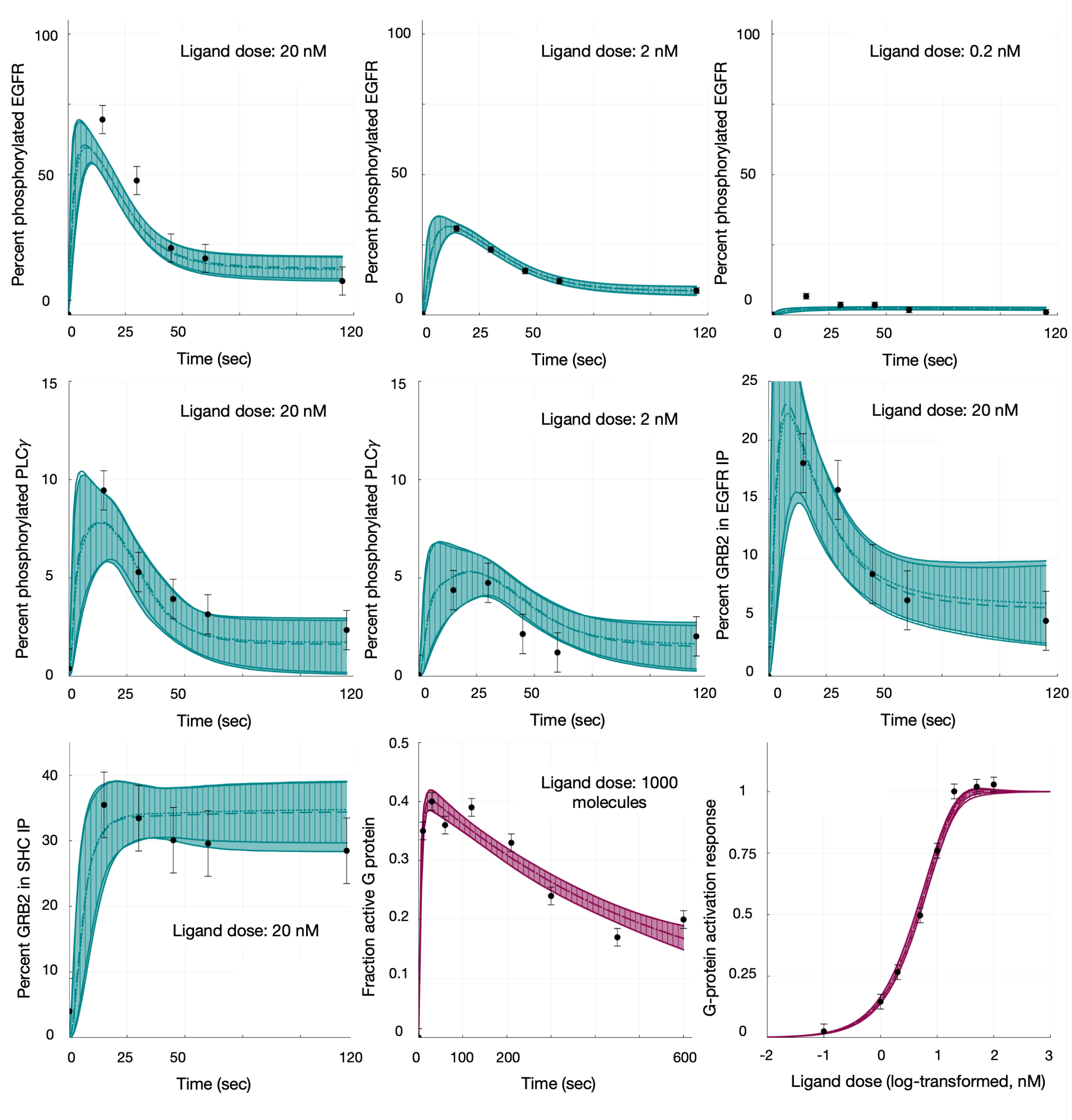


**Figure S3. Training data performance.** Cyan, EGFR; pink, GPCR. Shaded region, 90% quantiles of baseline approach; patterned region, 90% quantiles of augmented approach; dotted line, median prediction of augmented approach; dashed line, median prediction of augmented approach; black dots, experimental data with reported error.

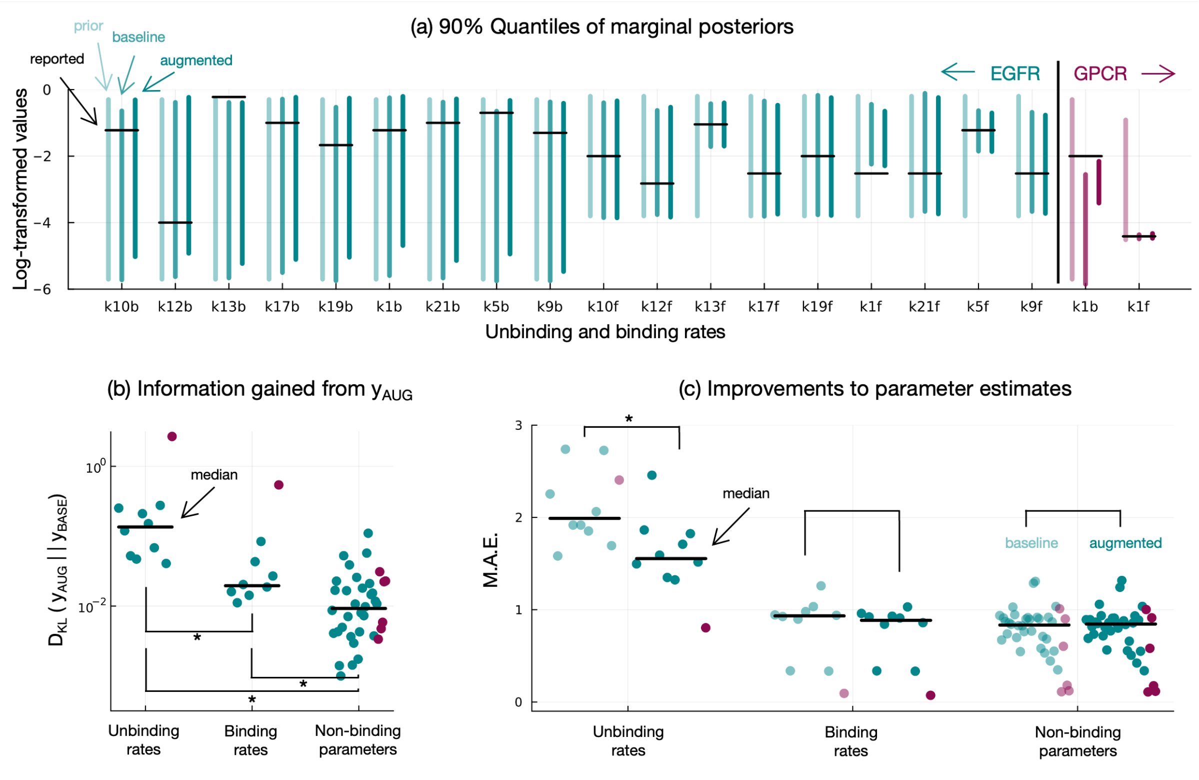


**Figure S4. Impact of considering Alpha Fold 3 confidence on parameter inference.** Cyan, EGFR results; pink, GPCR results. *p-value < 0.05. (a) 90% quantiles of marginal posterior distributions of binding parameters. All samples on log10 scale. Light cyan line, prior; medium cyan line, baseline posterior; dark cyan line, augmented posterior; black horizontal line, reported parameter value. (b) KL divergence, in bits, from baseline posterior to augmented posterior. Values are grouped by parameter function. (c) Mean absolute error (M.A.E.) of parameter samples. Mean taken with respect to each posterior distribution. Error calculated with respect to the values reported in the literature. Light cyan points, baseline; dark cyan points, augmented; black horizontal line, median M.A.E.

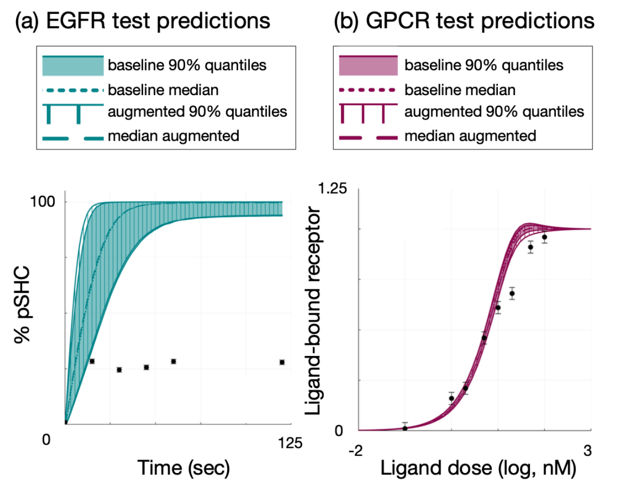


**Figure S5. Impact of considering AlphaFold 3 confidence on test predictions**. (a) Predictions for EGFR test set, the percent of phosphorylated signaling protein SHC from 0-120 seconds. (b) Predictions for GPCR test set, the amount of ligand bound receptor 60 seconds post-stimulation, at different ligand doses, relative to 1000 nM of ligand. Cyan, EGFR; pink, GPCR. Shaded region, 90% quantiles of baseline approach; patterned region, 90% quantiles of augmented approach; dotted line, median prediction of augmented approach; dashed line, median prediction of augmented approach; black dots, experimental data with reported error.

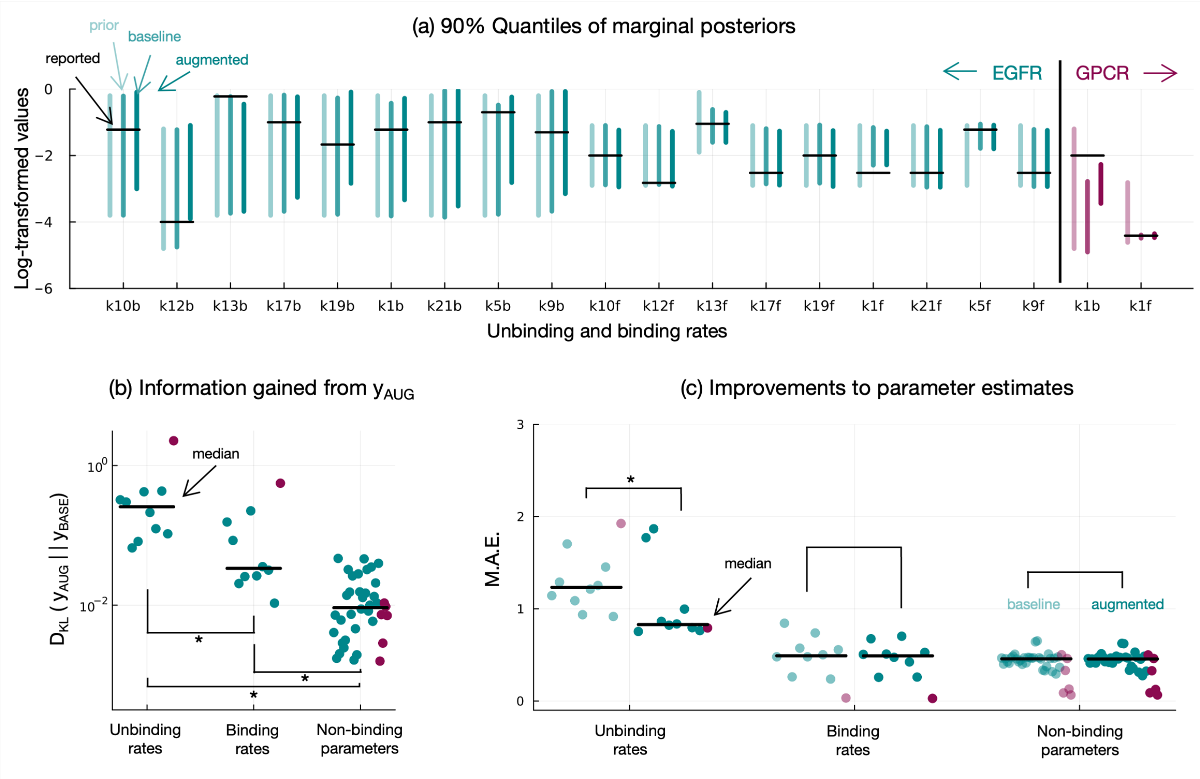


**Figure S6. Impact of more informative prior on parameter inference.** Cyan, EGFR results; pink, GPCR results. *p-value < 0.05 (a) 90% quantiles of marginal posterior distributions of binding parameters. All samples on log10 scale. Light cyan line, prior; medium cyan line, baseline posterior; dark cyan line, augmented posterior; black horizontal line, reported parameter value. (b) KL divergence, in bits, from baseline posterior to augmented posterior. Values are grouped by parameter function. (c) Mean absolute error (M.A.E.) of parameter samples. Mean taken with respect to each posterior distribution. Error calculated with respect to the values reported in the literature. Light cyan points, baseline; dark cyan points, augmented; black horizontal line, median M.A.E.

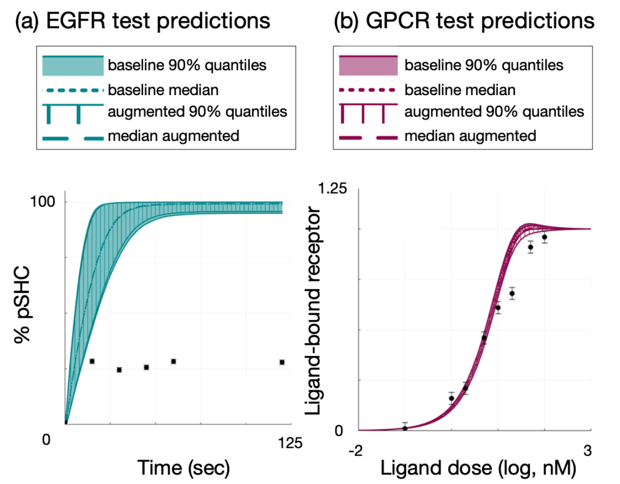


**Figure S7. Impact of more informative prior on test predictions**. (a) Predictions for EGFR test set, the percent of phosphorylated signaling protein SHC from 0-120 seconds. (b) Predictions for GPCR test set, the amount of ligand bound receptor 60 seconds post-stimulation, at different ligand doses, relative to 1000 nM of ligand. Cyan, EGFR; pink, GPCR. Shaded region, 90% quantiles of baseline approach; patterned region, 90% quantiles of augmented approach; dotted line, median prediction of augmented approach; dashed line, median prediction of augmented approach; black dots, experimental data with reported error.

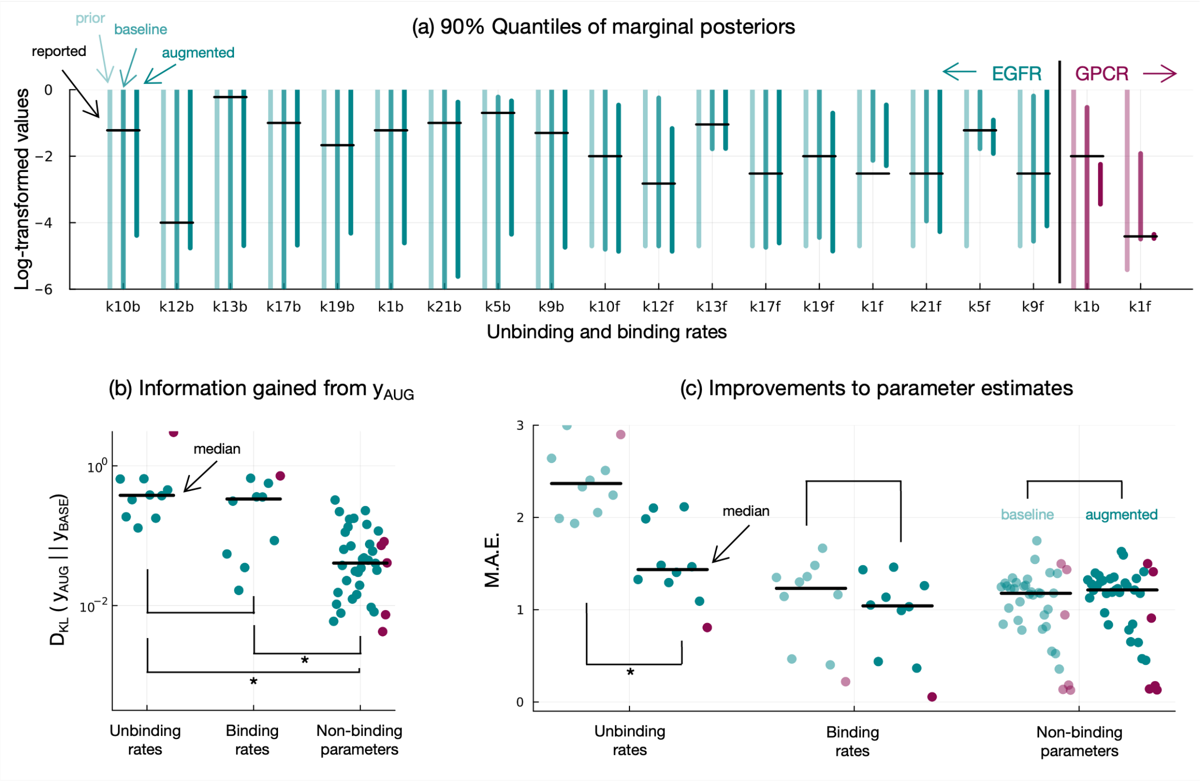


**Figure S8. Impact of less informative prior on parameter inference.** Cyan, EGFR results; pink, GPCR results. *p-value < 0.05 (a) 90% quantiles of marginal posterior distributions of binding parameters. All samples on log10 scale. Light cyan line, prior; medium cyan line, baseline posterior; dark cyan line, augmented posterior; black horizontal line, reported parameter value. (b) KL divergence, in bits, from baseline posterior to augmented posterior. Values are grouped by parameter function. (c) Mean absolute error (M.A.E.) of parameter samples. Mean taken with respect to each posterior distribution. Error calculated with respect to the values reported in the literature. Light cyan points, baseline; dark cyan points, augmented; black horizontal line, median M.A.E.

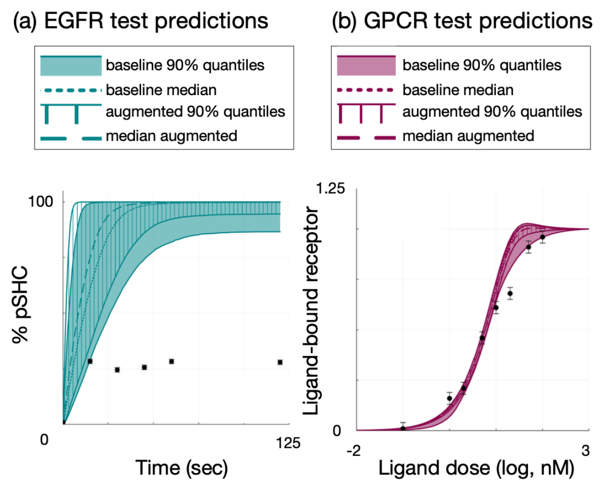


**Figure S9. Impact of less informative prior on test predictions**. (a) Predictions for EGFR test set, the percent of phosphorylated signaling protein SHC from 0-120 seconds. (b) Predictions for GPCR test set, the amount of ligand bound receptor 60 seconds post-stimulation, at different ligand doses, relative to 1000 nM of ligand. Cyan, EGFR; pink, GPCR. Shaded region, 90% quantiles of baseline approach; patterned region, 90% quantiles of augmented approach; dotted line, median prediction of augmented approach; dashed line, median prediction of augmented approach; black dots, experimental data with reported error.
